# Induction of plant disease resistance by mixed oligosaccharide elicitors prepared from plant cell wall and crustacean shells

**DOI:** 10.1101/2023.07.06.547984

**Authors:** Sreynich Pring, Hiroaki Kato, Sayaka Imano, Maurizio Camagna, Aiko Tanaka, Hisashi Kimoto, Pengru Chen, Abhijit Shrotri, Hirokazu Kobayashi, Atsushi Fukuoka, Makoto Saito, Takamasa Suzuki, Ryohei Terauchi, Ikuo Sato, Sotaro Chiba, Daigo Takemoto

## Abstract

Basal plant immune responses are activated by the recognition of the conserved pathogen-associated molecular patterns (PAMPs), or breakdown molecules released from plants after damage by pathogen infection, so-called danger-associated molecular patterns (DAMPs). While chitin-oligosaccharide (CHOS), a primary component of the fungal cell wall, is most known as PAMP, plant cell wall-derived oligosaccharides, cello-oligosaccharides (COS) from cellulose and xylo-oligosaccharide (XOS) from hemicellulose, are representative DAMPs, which activate signaling steps similar to PAMP-induced immunity to elicit defenses and provide protection against pathogens. In this study, elicitor activities of COS prepared from cotton linters, XOS prepared from corn cobs as well as chitin-oligosaccharide (CHOS) from crustacean shells were comparatively investigated. In Arabidopsis, treatment of COS, XOS or CHOS triggered typical defense responses such as reactive oxygen species (ROS) production, activation of MAP kinases phosphorylation, callose depositions, and activation of the promoter for defense-related transcription factor *WRKY33*. When COS, XOS and CHOS were used at concentrations with similar activity in inducing ROS production and callose depositions, CHOS was particularly highly potent in activating the MAPK kinases and *WRKY33* promoters. Among the COS and XOS with different degrees of polymerization (DP), cellotriose (DP3) and xylotetraose (DP4) showed the highest activity for the activation of *WRKY33* promoter. Simultaneous treatment of COS, XOS and CHOS leads to a strong transcriptional change for defense-related genes, and gene ontology (GO) enrichment analysis of RNAseq data revealed that a mixture of three oligosaccharide (oligo-mix) effectively activate the plants disease resistance. In practice, treatment of the oligo-mix enhanced the resistance of tomato to powdery mildew, but plant growth was not inhibited but rather tended to be promoted, providing evidence that mixed oligosaccharides have beneficial effects on improving disease resistance in plants, making them a promising class of compounds for practical application.

## INTRODUCTION

Plants are constantly confronted with a wide variety of pathogens, including fungi, oomycetes, and bacteria. To counteract those pathogens, plants have evolved an intricate immune system, which can recognize conserved pathogen- or microbe-associated molecular patterns, known as PAMPs/MAMPs, including epitopes of bacterial flagella, lipopolysaccharides, peptidoglycans, fungal chitin and fungal/oomycete β-glucan and sphingolipids (Monaghan and Zipfel, 2012; Kato et al., 2022; Ngou et al., 2022). Aside from PAMP perception, plants are also able to detect endogenous signals of cell damages, such as extracellular ATP, oligogalacturonides and oligosaccharides derived from plant cell wall, which are released during pathogen infection, insect attack or other stress conditions, so-called damage-associated molecular patterns (DAMPs) (Choi and Klessig, 2016). Receptor-mediated recognition of PAMPs and DAMPs at the cell surface is an innate immune mechanism common to plants, animals, and insects (Jones and Takemoto, 2001; Boller and Felix, 2009; Tang et al., 2012; Ngou et al., 2022; Zhao et al., 2023).

The plant cell wall is the critical site of initial encounters between plants and their microbial pathogens. A plant cell wall is mainly composed of high molecular weight polysaccharides such as cellulose, hemicellulose, pectin, and lignin, which are complex and dynamic polysaccharide structures to protect plants from their surrounding environments (Somerville et al., 2004). Cell wall heterogeneity has evolutionary implications for the different mechanisms by which pathogens evolved to breach plant cell walls, including the secretion of a variety of cell wall-degrading enzymes (CWDEs) such as cellulases, polygalacturonases, or xylanases (Kubicek et al., 2014). Monitoring systems for cell wall integrity respond to cell wall restructuring caused by pathogen infection, abiotic stress, and cell expansion during growth and development, triggering compensatory responses. Changes in cell wall composition or integrity caused by physical or chemical means have significant impacts on plant resistance to pathogens and abiotic stresses because they typically activate defensive signaling pathways, some of which are regulated by plant hormones (Hou et al. 2019).

Cellulose is the central morphogenic polysaccharide to stabilize the plant cell wall. It is synthesized by the cellulose synthase complex, which converts UDP-Glc into β-1,4-glucan chains in the cell wall to form cellulose microfibrils. Cello-oligosaccharides (COS) are short-chain linear polymers that can be produced by partial hydrolysis of cellulose (Chen et al., 2019). COS and mixed-linked glucans (β-1,4/β-1,3-glucans) has been reported as elicitors to activate defense responses to increase disease resistance (de Azevedo Souza et al., 2017; Johnson et al., 2018; Barghahn et al., 2021; Rebaque et al., 2021; Zarattini et al., 2021).

Hemicellulose consists of heterogeneous polysaccharides, and its most important biological role is to interact with cellulose and lignin, contributing to the strengthening of cell walls (Scheller and Ulvskov, 2010). Xylo-oligosaccharide (XOS) is a major component of hemicellulose, which is made up of xylose monomeric units bound together by β-1,4 linkages. XOS are found naturally in fruits, vegetables, milk, and honey, and can be produced commercially through the enzymatic, chemo-enzymatic, and chemical hydrolysis of xylan in hemicellulose from a variety of sources including plant materials (Chen et al., 2021). XOS has been reported to have elicitor activity as a DAMP in several plant species. Although resistance induction is generally considered to be a trade-off for growth, previous report indicated that XOS can promote plant growth and development (Cutillas-Iturralde and Lorences, 1997; Kaida et al., 2010).

Oligosaccharides, which have been reported to promote disease resistance and plant growth, have the potential to be used effectively in the field as so-called biostimulants. Firstly, in this study, elicitor activity of DAMPs derived from the plant cell wall, COS and XOS as well as a representative PAMP, CHOS was comparatively investigated on Arabidopsis by measuring typical defense responses such as ROS production, MAP kinase activation, callose deposition and activation of defense related *WRKY33* promoter. By employing RNAseq followed by GO enrichment analysis, overall effects of single and simultaneous treatments of these oligosaccharides (oligo-mix) on treated plants were analyzed. For COS and XOS, the degree of polymerization with higher elicitor activity was also determined. Moreover, the effects of oligo-mix treatment on tomatoes grown in greenhouses on disease resistance and on the growth were analyzed to evaluate the efficacy of an oligo-mix as a biostimulant.

## MATERIALS AND METHODS

### Plant Material and Growth Conditions

Seed of *Arabidopsis thaliana* Col-0 or transgenic line containing luciferase reporter under the control of *WRKY33* (AT2G38470) promoter (Kato et al., 2022) were used in this study. Seeds were surface sterilized with 3% hydrogen peroxide, 50% EtOH solution by gentle shaking for one minute. Subsequently, the sterilization solution was replaced with sterilized water, then individual seeds were grown in separate wells of a 96-micro well plate (Nunc 96F microwell white polystyrene plate, Thermo Fisher Scientific, Waltham, MA, United States) containing 150 μl of Murashige and Skoog (MS) liquid medium (1/2 MS salts, 0.05% [w/v] MES, 0.5% [w/v] Sucrose, adjusted to pH 5.8 with NaOH) and covered with a clear plastic cover. For luciferase-mediated promoter analysis, MS liquid medium was supplemented with 50 μM D-Luciferin potassium salt (Biosynth Carbosynth, Compton, UK). The plate was placed in a controlled growth incubator for 12 days at 23 °C with 24 h light.

### Elicitors Used in This Study

Cello-oligosaccharide were prepared by hydrolysis of cotton linters or microcrystalline cellulose (Avicell PH 101, Sigma-Aldrich, Burlington, MA, USA) (Chen et al., 2019), and chitin-oligosaccharide from crab and shrimp shells (Kobayashi et al., 2017) as previously reported. Xylo-oligosaccharide prepared from corn cob was provide from Resonac Corporation (former Showa Denko K. K., Tokyo, Japan). Flg22 peptide was purchased from GenScript Biotech Corporation (Piscataway, NJ, USA).

### Measurement of Reactive Oxygen Species (ROS) Production

The relative intensity of ROS generation was determined by counting photons from L-012-mediated chemiluminescence as reported previously (Imano et al., 2022). Arabidopsis seedlings were grown in MS medium in separate wells of a 96-micro well plate as described above for 12 days, and MS medium was replaced with 100 μl distilled water one day before the measurement. The water was then replaced with 100 μl of reaction solution containing 1 mM L-012 (Fujifilm Wako Pure Chemical, Osaka, Japan) and indicated elicitor(s). Chemiluminescence was measured by a multiplate reader (TriStar LB941; D-75323 Berthold Technologies, Bad Wildbad, Germany).

### MAPK Activation Assay

MAPK activation assays were performed as previously reported (Kato et al., 2022). Arabidopsis seedlings were grown in MS liquid medium as describe above. Seedlings were then elicited with indicated elicitors for 15 or 30 min and frozen in liquid nitrogen. Proteins were extracted in extraction buffer (50 mM HEPES-KOH pH 7.4, 5 mM EDTA, 0.5 mM EGTA, 50 mM β-glycerophosphate, 10 mM NaF, 10 mM Na_3_VO_4_, and 2 mM DTT). MAPK activation was detected by western blot with Phospho-p44/42 MAPK (Erk1/2; Thr-202/Tyr-204) rabbit monoclonal antibodies #9101 (Cell Signaling Technology, Danvers, MA, USA). Blots were stained with PageBlue Protein Staining Solution (Thermo Fisher Scientific, Waltham, MA, USA) to verify equal loading.

### Staining of Callose Deposition

Staining of callose deposition was conducted as previously reported (Shibata et al., 2016). Arabidopsis seedlings grown in MS liquid medium were treated with indicated elicitors for 24 h, and seedlings were fixed in fixation solution (1% [v/v] glutaraldehyde, 5 mM citric acid, and 90 mM Na_2_HPO_4_, pH 7.4) overnight. Fixed seedlings were then incubated in 1 ml hot water at 100 °C for 5 minutes, then decolorized with 99% EtOH, and stained in aniline blue stain solution (0.1% (w/v) water-soluble aniline blue in 67 mM phosphate buffer, pH 12.0) to detect callose deposition. The fluorescence spots of deposited callose was detected by a BX51 fluorescence microscope (Olympus, Tokyo, Japan) using an excitation wavelength of 365 nm. The number of callose spot per area was quantified in ImageJ software (Schneider et al., 2012). Data represents mean ± SD (n = 5, leaves from five seedlings).

### Measurement of Promoter Activity of *WRKY33* gene

Transgenic Arabidopsis containing *Luciferase* marker gene under the control of defense related *WRKY33* promoter (Kato et al., 2022) were grown as described above. Bioluminescence intensity was monitored from 12-day-old seedlings after the treatment with water- or indicated elicitors using a multiplate reader TriStar LB941 (Berthold Technologies, Bad Wildbad, Germany) (Kato et al., 2020). Data are shown as means ± SE (n = 8 seedlings) from three independent repetition with similar results.

### RNAseq/Gene Ontology Enrichment Analysis of Arabidopsis Treated with Oligosaccharides

RNAseq analysis was principally performed as previously described (Imano et al., 2022). Total RNA was isolated from Arabidopsis seedlings (8 seedlings per sample) using the RNeasy Plant Mini Kit (QIAGEN, Hilden, Germany) 24 h after the elicitor treatment. Libraries were constructed using KAPA mRNA Capture Kit (Roche Diagnostics, Tokyo, Japan) and MGIEasy RNA Directional Library Prep Set (MGI, Shenzhen, China), and sequenced on DNBSEQ-G400RS (MGI) with 150 bp paired-end protocol. The RNA-seq reads were filtered using trim-galore v.0.6.6 (Martin, 2011, bioinformatics.babraham.ac.uk) and mapped to the *Arabidopsis thaliana* genome (genome assembly TAIR10.1, NCBI RefSeq sequence GCF_000001735.4) using HISAT2 v.2.2.1 (Kim et al., 2019) and abundance inferred via StringTie v.2.1.7 (Kovaka et al., 2019). Significant differential expression was determined using DESeq2 v.1.32.0 (Love et al., 2014). All software used during RNA-seq analysis was run with default settings. Gene ontology (GO) enrichment analysis was performed using PANTHER statistical overrepresentation test (http://pantherdb.org; Version 17.0, Thomas et al. 2022) using default settings (Fisher’s exact test, False discovery rate (FDR) < 0.05). Heatmap was plotted using SRplot (https://www.bioinformatics.com.cn/en), a free online platform for data analysis and visualization. RNA-seq data for Arabidopsis reported in this work are available in GenBank under the accession numbers DRA016697.

### Fractionation of Oligosaccharides

The separation of individual cello-oligosaccharides from the hydrolysis product solution was performed by using high performance liquid chromatography (HPLC) equipped with a refractive index detector (Shimadzu RID 10-ATVP, Shimadzu, Kyoto, Japan) and a fraction collector (Shimadzu FRC 10A, Shimadzu). Three Shodex OHpak SB-802.5 HQ columns (Resonac, Tokyo, Japan) in series (ø 8 × 300 mm; eluent, water at 0.5 ml/min; 55°C) were used to separate cello-oligosaccharides up to cellohexaose. Individual fractions were collected based on retention time observed in chromatogram. 50 μl of sample was injected for one fraction collection run and the process was repeated 100 times. The separated cello-oligosaccharides solutions were collected in an eggplant-shaped flask and water was removed using a rotary evaporator. The obtained sample was redissolved in 5 ml water and quantified. The volume of solution was adjusted to obtain a 50 ppm solution. The separation of xylo-oligosaccharide and chitin-oligosaccharide was done by following the same method by using three Shodex GF-210 (HQ) HPLC columns (Resonac) in series (ø 8 × 300mm; eluent, water at 0.5 mL min^−1^; 50°C).

### Inoculation Assay of Tomato

Tomato plants (cvs. Frutica and Momotaro) were grown in a greenhouse at Fukui prefecture, Japan from April to July 2020. After germination of seedlings, plants were sprayed with water or oligo-mix (20 mg/ml COX, 40 mg/ml XOS and 20 mg/ml CHOS), once every two weeks. Plants were grown and naturally allowed to get infected with powdery mildew in the greenhouse. After 6 spraying treatments (approx. 3 months after potting), disease symptoms which had appeared on the leaves (the number of colonies) were quantified from 6 plants for each treatment per experiment. Data are shown as mean ± SE.

### RNAseq Analysis of Tomato Treated with Oligo-mix

Tomato plants (cv. Renaissance) were grown in a growth room at 23°C with 16 h of light per day. After germination of seedlings, plants were sprayed with water or oligo-mix (20 mg/ml COX, 40 mg/ml XOS and 20 mg/ml CHOS) from 3 days after the seedling germination and then once every week (for 4 times). One week after the last treatment, growth of tomato was measured, and the leaf samples were used for the extraction of total RNA using the RNeasy Plant Mini Kit (QIAGEN). Evaluation of RNA quality, construction of library and sequencing were performed as previously described (Rin et al., 2021; Kuroyanagi et al., 2022). Filtering of reads, and mapping to tomato genome (ver SL4.0, https://solgenomics.net/ftp/tomato_genome/, Hosmani et al., 2019) were performed as described above. RNA-seq data for tomato reported in this work are available in GenBank under the accession numbers DRA016704.

## RESULTS

### Oligosaccharide Elicitors Induce Immune Responses in *Arabidopsis* seedling

In this study, cello-oligosaccharide (COS) prepared from cotton linters, xylo-oligosaccharide (XOS) prepared from corn cobs, and chitin-oligosaccharide (CHOS) from shrimp shells were prepared (Kobayashi et al., 2017, Chen et al., 2019) and examined on their ability to induce plant defense responses. These oligosaccharide elicitors included mixed degrees of polymerization (DP). The CHOS solution included monomeric N-acetylglucosamine, dimers (DP2) up to nonamer (DP9) chitins, COS solution included monomeric glucose and DP2 to DP9 cello-oligosaccharides, and XOS solution contained DP2 to hexameric (DP6) xylo-oligosaccharides (Figure 1A). Elicitor activity of these oligosaccharides was investigated on seedlings of the model plant Arabidopsis. During preliminary test using 20 mg/ml oligosaccharide elicitors, XOS showed significantly weaker elicitor activity compared with the other oligosaccharides, thus 40 mg/ml XOS was used in the following experiments.

**FIGURE 1.**
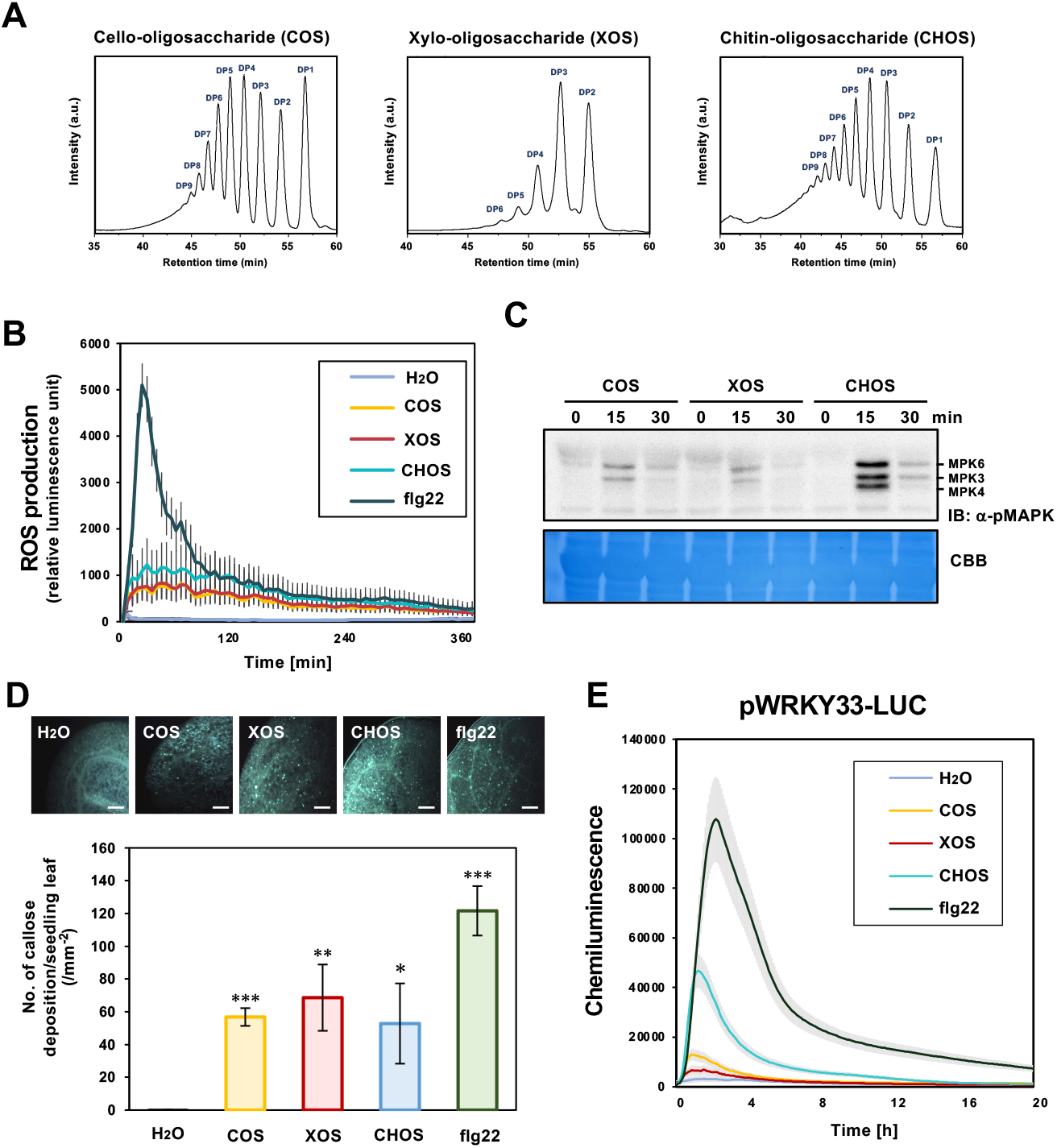
Induction of defense responses in Arabidopsis seedlings treated with oligosaccharides derived from plant cell wall and crustacean shell. **(A)** HPLC chromatogram of cellooliogaccharide (COS) prepared from cotton linters, xylooligosaccharide (XOS) prepared from corn cobs, and chitinoligosaccharide (CHOS) prepared from crustacean shells to analyze the distribution of degree of polymerization (DP). **(B)** Reactive oxygen species (ROS) production was detected in Arabidopsis seedling after treatment with 20 mg/ml COS, 40 mg/ml XOS, 20 mg/ml CHOS or 100 nM flg22. Data are means ± standard error (n = 8). **(C)** MAP kinase activation in Arabidopsis seedlings after treatment with 20 mg/ml COS, 40 mg/ml XOS or 20 mg/ml CHOS. The phosphorylation of MAP kinases was determined by western blot using phospho-p44/42 MAPK (Erk1/2; Thr-202/Tyr-204) antibody. **(D)** Arabidopsis seedlings were treated with indicated elicitors as (A) and callose deposition was counted by aniline blue staining at 24 h after treatments. Fluorescence spots were counted from five leaves of five individual seedlings (n = 5) for each treatment and detected by fluorescence microscopy. Bars = 100 μm. **(E)** Activation of *AtWRKY33* was detected as chemiluminescence in Arabidopsis transformant pWRKY33-LUC treated with oligosaccharide elicitors in (A). All data marked with asterisks are significantly different from H_2_O as determined by the two-tailed Student’s *t* test. ***p<0.001, **p< 0.01, *p< 0.05.

Production of reactive oxygen species (ROS) and activation of MAP kinases are common reactions which are initially activated during the induction of plant disease resistance (Doke, 1983, Torres et al., 2002, Xu et al., 2016). After the treatment of Arabidopsis seedlings with oligosaccharide elicitors, ROS production was activated within 5 min and peaked at approx. 30 min before decreasing (Figure 1 B). More intense activation of ROS production was observed with 100 nM flg22 peptide (approx. 227 ng/ml), a representative bacterial PAMP, around 2 h after the treatment, although continuous ROS production after 2 h did not differ significantly among all elicitors. Activation/phosphorylation of defense-related MAP kinases, MPK6 and MPK3 (Asai et al., 2002), were induced within 15 min after the treatment with COS, XOS and CHOS, and then attenuated at 30 min. Significant activation of MPK6 and MPK3 was detected by CHOS treatment, compared with that by other oligosaccharides, and clear activation of MPK4 (Ichimura et al., 2006) was only detected after the treatment with CHOS (Figure 1 C). Another typical defense response, deployed to reinforce plant cell walls during a pathogens attack, is callose deposition (Ellinger et al., 2014). After treatment of Arabidopsis seedlings with COS, XOS, CHOS or flg22 for 24 h, marked increase in callose depositions was observed (Figure 1D), but in contrast to the MAP kinase activation, the three oligosaccharides were not significantly different regarding callose deposition. The Arabidopsis transcription factor WRKY33 is a central regulator for the regulation of defense-related genes during the attack of pathogens (Zheng et al., 2006, Birkenbihl et al., 2012). Transgenic Arabidopsis with luciferase (*Luc*) reporter gene under the control of the *WRKY33* promoter (pWRKY33-LUC, Kato et al., 2022) was used to detect the activity of the *WRKY33* promoter after oligosaccharide treatment. Activation of the *WRKY33* promoter was detected within 20 min after the treatment with COS, XOS or CHOS and peaked at approx. 1 h after treatment (Figure 1E), and CHOS showed significantly stronger activity compared with COS and XOS. These results confirm that COS, XOS and CHOS treatments activate typical plant defense responses, with CHOS being particularly active in inducing MAPK phosphorylation and *WRKY33* promoter activation compared to the other oligosaccharides. There was no clear difference in activity among the three oligosaccharides for ROS generation and callose accumulation. Recently, pattern recognition receptors (and potential co-receptors) for COS recognition have been isolated (namely CORK1/AT1G56145, AT1G56130 and AT1G56140), which are distinct from CERK1 and LYK5-mediated CHOS recognition mechanisms (Miya et al., 2007; Cao et al., 2014; Tseng et al., 2022; Martín-Dacal et al., 2023). It is however still not clear how distinct the downstream signaling factors of these PAMP/DAMP recognitions are, given their similar defense response outcomes.

### Expression Profiles of Arabidopsis Genes after the Treatment with Cello-, Xylo-, Chitin-oligosaccharide or Their Mixture

To examine the expression profiles of Arabidopsis genes induced by oligosaccharide elicitors, RNA-seq analysis was performed on Arabidopsis seedlings 24 h after oligosaccharide treatment. Since the mixture of all three oligosaccharides has an addictive effect on the activation of *WRKY33* promoter (Figure 2A), RNA-seq analysis of seedlings treated with the mixture (oligo-mix) was also performed. A large number of genes (287 for COS, 216 for XOS and 193 for CHOS, respectively) was significantly up-regulated (log_2_-fold change value > 2, P < 0.05, average TPM > 1) in Arabidopsis treated with oligosaccharide elicitors, and 126 genes were commonly induced by all three oligosaccharides (Figure 2B). Among the three oligosaccharide elicitors, COS and XOS shared the highest commonly induced genes (193 genes). Consistently, hierarchical clustering of significantly up- or down-regulated genes revealed that treatment with COS and XOS elicits similar gene expression patterns in Arabidopsis seedling (Figure 2C).

**FIGURE 2.**
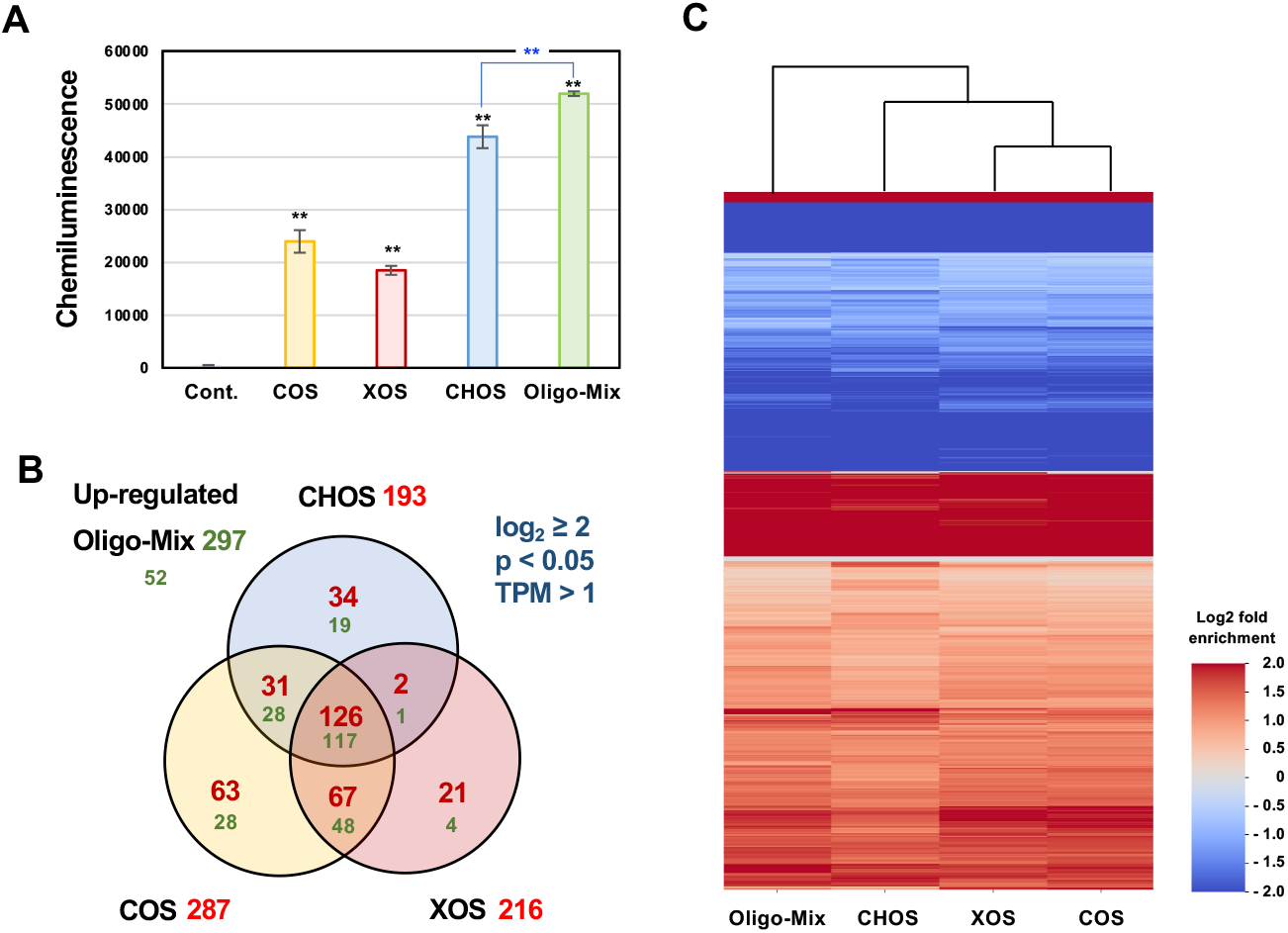
RNA-seq analysis of Arabidopsis treated with oligosaccharide elicitors or their mixture. **(A)** Arabidopsis transformant pWRKY33-LUC containing *LUC* transgene under the control of *AtRKY33* promoter was treated with 20 mg/ml cello-oligosaccharide (COS), 40 mg/ml xylo-oligosaccharide (XOS), 20 mg/ml chitin-oligosaccharide (CHOS), and their mixture (Oligo-mix). Total chemiluminescence for 12 h after the elicitor treatment was shown. Data marked with asterisks are significantly different compared with control (or between samples, blue asterisks) as determined by the two-tailed Student’s *t* test. **p<0.01. **(B)** Venn diagram representing the up-regulated differentially expressed genes (DEGs) in Arabidopsis treated with COS, XOS, CHOS and mixture treatment for 24 h, which are selected based on TPM > 1, log_2_ > 2 and p<0.05. Green numbers indicate the number of up-regulated genes induced by oligo-mix. **(C)** Heatmap analysis of differentially expressed genes for the oligosaccharide treatments. The color bar represents the fold change value (log2) with p<0.05.

Although genes up-regulated by treatment with the mixture of the three oligosaccharides (oligo-mix) were similar to those upregulated by treatment with single oligosaccharides, expression of some genes were enhanced by treatment with oligo-mix. For example, genes encoding enzymes for production of camalexin (the major phytoalexin of Arabidopsis, Schuhegger et al., 2006; Nafisi et al., 2007), namely AT2G30750 (*CYP71A12*), AT2G30770 (*CYP71A13*) and AT3G26830 (*PAD3*) were significantly upregulated by oligo-mix (Figure 3). Genes induced by oligosaccharides with high TPM values (>100) contained some disease resistance-related genes, including plant defensins *PDF1*.*1* and *1*.*4* (AT1G75830 and AT1G19610, Thomma et al., 2002), chitinase *PR-3 like* (AT2G43590, Passarinho et al., 2001), *NATA1* for production of defensive metabolite N^δ^-acetylornithine (AT2G39030, Adio et al., 2011), *SWEET2* encoding vacuolar sugar transporter to reduce sugar availability for pathogen (AT3G14770, Chen et al., 2015) (Figure 4A). Several genes related to plant cell wall strengthening that may be involved in pre-invasion defense are also induced, such as cell-wall hydroxyproline-rich glycoprotein extensin *EXT1* (AT1G76930, Wei and Shirsat, 2006), *Prx37* and *CASPL1D1* probably involved in the formation of lignin-based barrier (AT4G08770 and AT4G15610, Pedreira et al., 2011; Lee at al., 2019) and pectin methylesterase PME17 (AT2G45220, Del Corpo et al., 2020) (Figure 4A). Genes involved in the response to oxidative stress, including glutathione S-transferases *GSTU24* and *GSTF7* (AT1G17170 and AT1G02920) and thioredoxin-dependent peroxidase *TPX2* (AT1G6597) were also significantly up-regulated (Figure 4B). In addition to that, several genes that are potentially related to plant root development were also induced such as root meristem growth factor *RGF7/GLV4* (AT3G02240, Fernandez et al., 2013), asparaginases *ASPGB1* (AT3G16150, Ivanov et al., 2012) and nitrilase *NIT2* (AT3G44300) (Figure 4C). While in some cases the oligo-mix treatment resulted in additive expression compared to the single treatment, in others the expression was not significantly different from the single treatment, or rather the expression was slightly suppressed by the oligo-mix treatment (Figures 3). These results suggest that the eventual phenotypes (i.e. expression level of each gene) are reflected by the interplay of multiple signaling in plant cells that simultaneously recognize multiple PAMPs and DAMPs.

**FIGURE 3.**
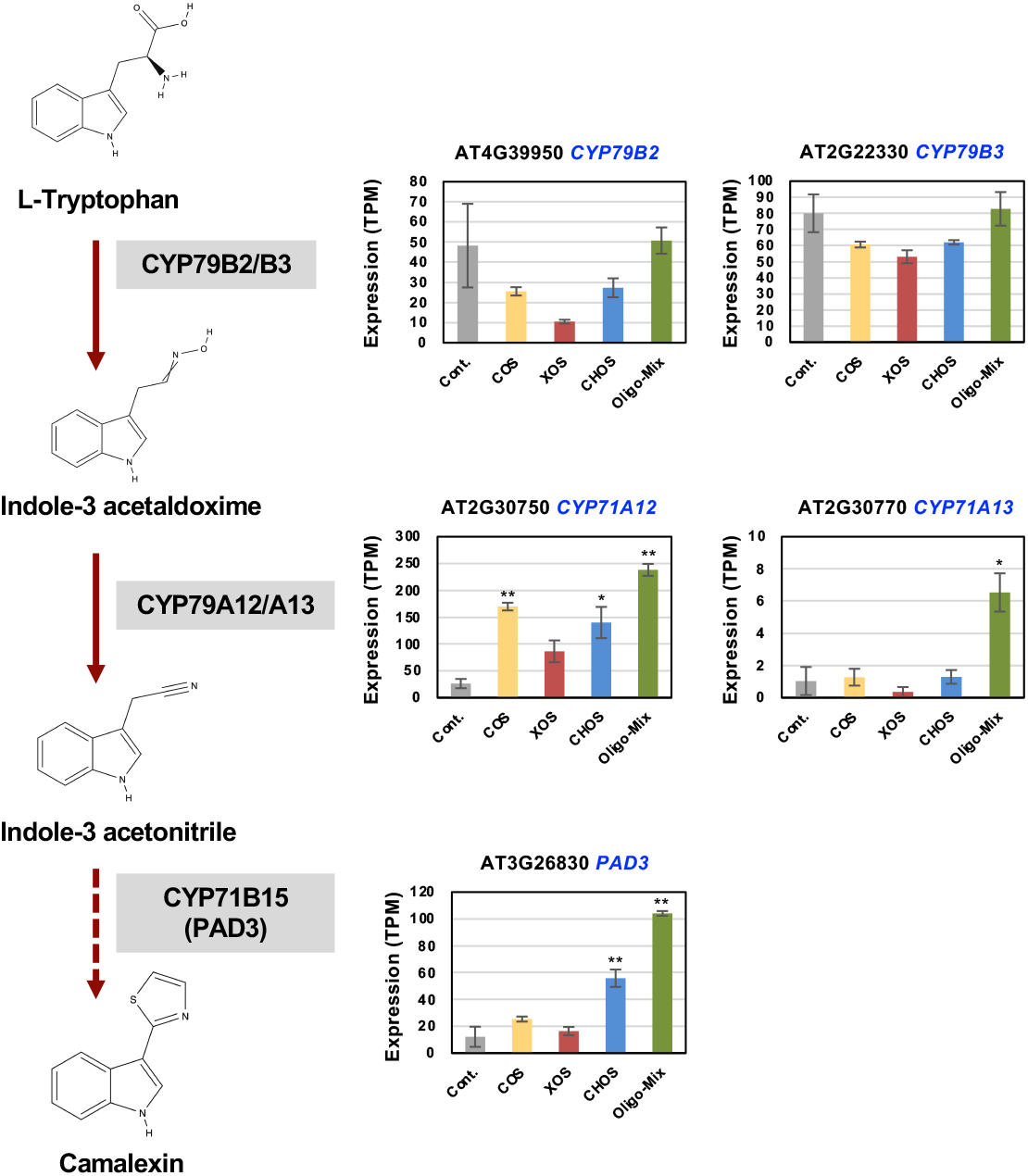
Expression profiles of Arabidopsis genes encoding enzymes involved in the camalexin biosynthesis. Gene expression (TPM value) was determined by RNA-seq analysis of Arabidopsis seedlings treated with 1% DMSO (Cont.), 20 mg/ml cello-oligosaccharide (COS), 40 mg/ml xylo-oligosaccharide (XOS), 20 mg/ml chitinoligosaccharide (CHOS) or the mixture for 24 h. Data are means ±SE (n = 3). Data marked with asterisks are significantly different from control as assessed by the two-tailed Student’s t test: **p<0.01, *p<0.05.

**FIGURE 4.**
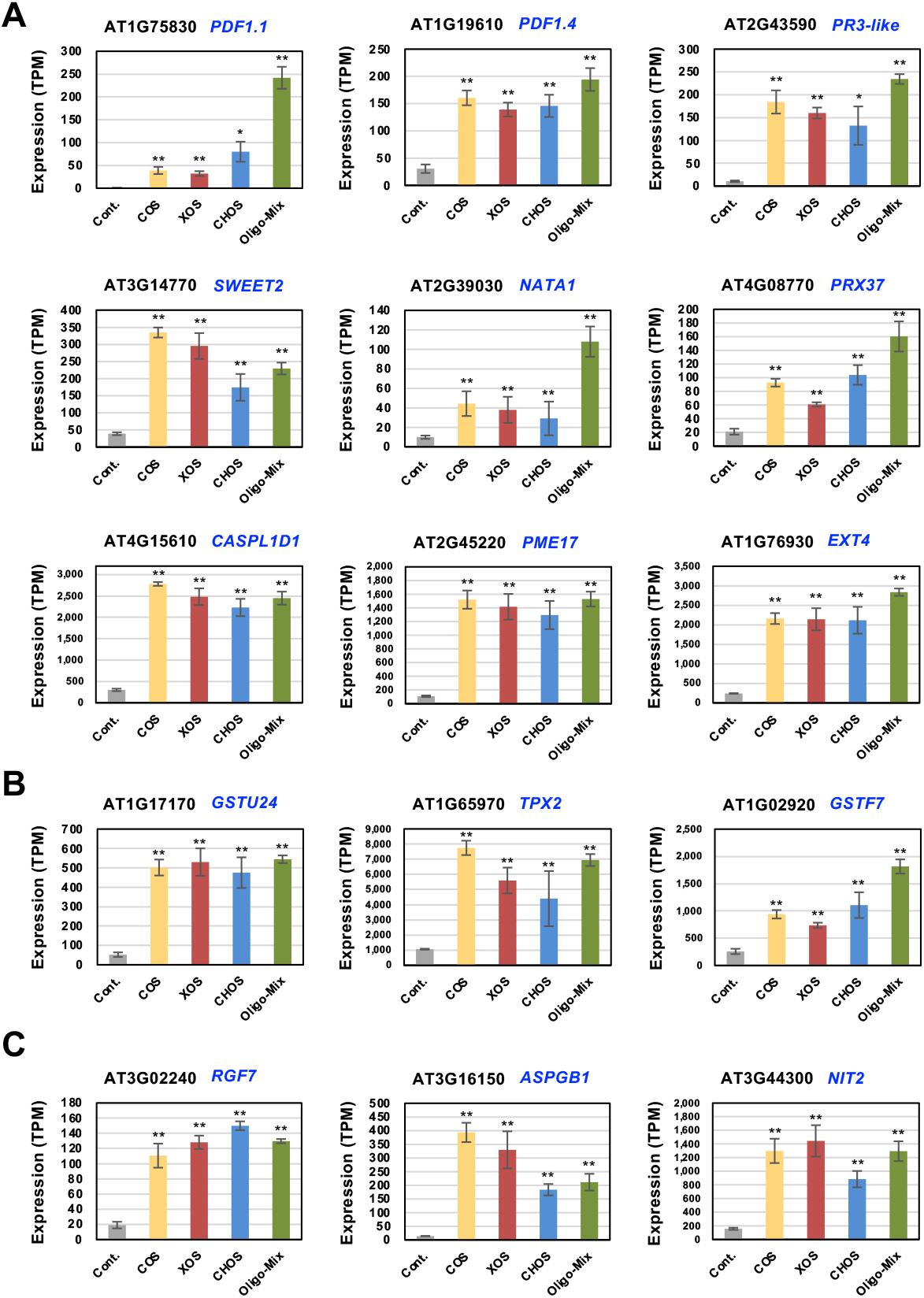
Representative Arabidopsis genes upregulated by treatment with oligosaccharides and their mixture. Gene expression (TPM value) was determined by RNA-seq analysis of Arabidopsis seedlings treated with 1% DMSO (Cont.), 20 mg/ml COS, 40 mg/ml XOS, 20 mg/ml CHOS or their mixture (Oligo-mix, 20 mg/ml COS, 40 mg/ml XOS, 20 mg/ml CHOS) for 24 h. **(A)** Genes related to plant disease resistance and cell wall reinforcement. Genes related to oxidative stress **(B)** and root development **(C)**. Data marked with asterisks are significantly upregulated from control as determined by two-tailed Student’s *t-*test **p<0.01, *p<0.05.

### Gene Ontology Enrichment Analysis for Oligosaccharide-induced Genes in Arabidopsis

To determine the overall influence of treatment with oligosaccharide elicitors on the plant, genes upregulated by oligosaccharide elicitors were assigned to gene ontology (GO) terms using the analysis tool PANTHER (Thomas et al., 2022) for GO enrichment analysis (Figure 5 and Supplementary Figures 1-3). This analysis confirmed that oligosaccharide elicitors and their mixture can enhance the biological processes (BP) categories related to plant defense against microorganisms such as defense responses to fungus (GO:00508320), defense responses to bacterium (GO: 0042742), response to oomycetes (GO:0002239) and systemic acquired resistance GO:0009627. Gene subsets associated with BP categories related to stress response, including response to stress (GO:0006950), toxin metabolic process (GO:0009404) and response to oxidative stress (GO:0006979), were also shown to be activated by oligosaccharide elicitors. The three oligosaccharides appear to share most pathways, but GO terms related to resistance to insects, defense response to insect (GO:0002213) and response to insect (GO:0009625), were enriched only by the chitin treatment (and oligo-mix). This result may be related to the fact that chitin is the major component of the insect exoskeleton (Zhu et al., 2016). Activated cellular component (CC) categories include extracellular region (GO:0005576) and apoplast (GO:0048046), implicating the activation of defense responses to external attack. Taken together, these results suggest that oligosaccharide-mediated gene expression is associated with (GO) terms which function in plant immune responses (Figure 5 and Supplementary Figures 1-3).

**FIGURE 5.**
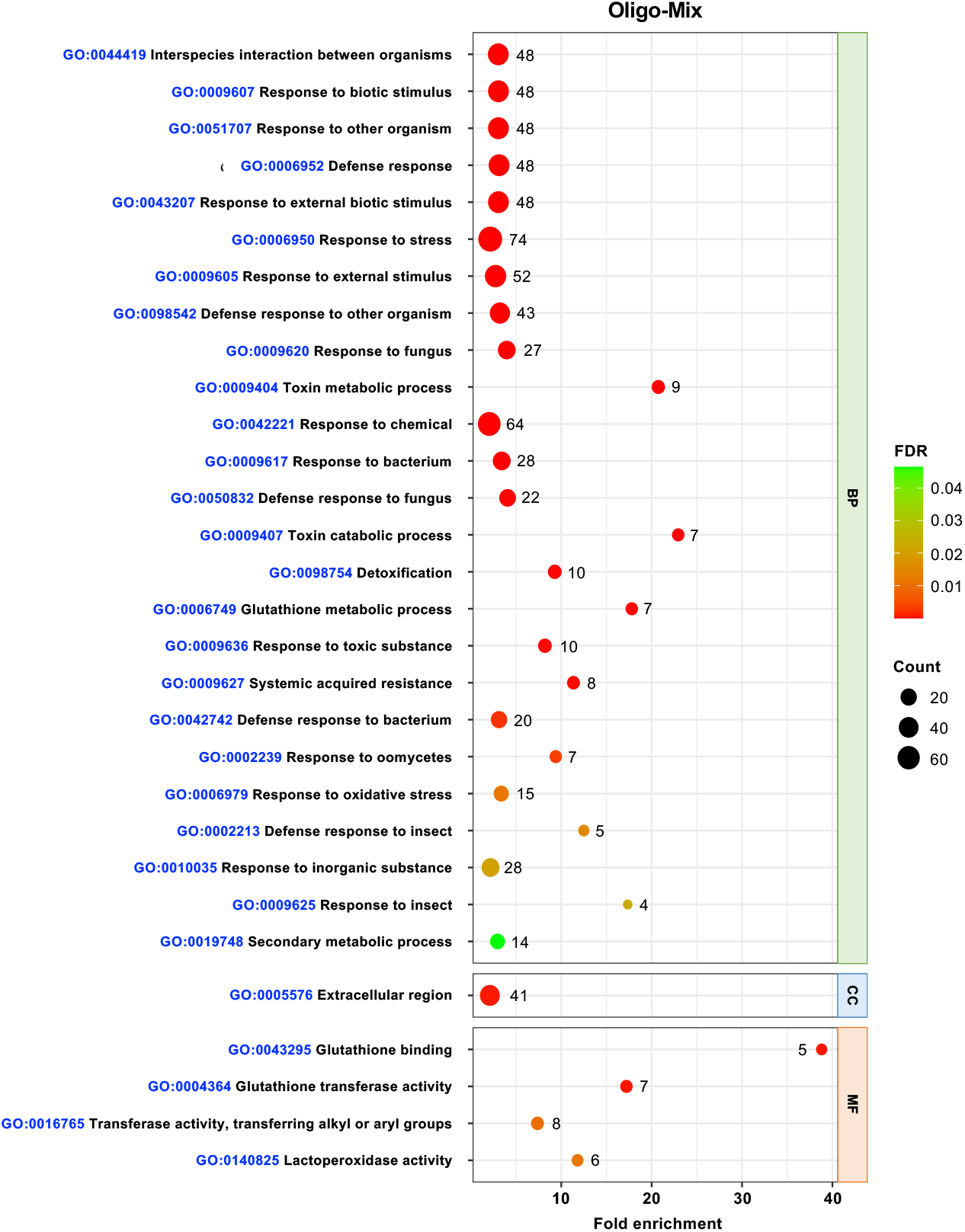
Gene ontology (GO) enrichment analysis of up-regulated genes in Arabidopsis treated with oligo-mix (20 mg/ml cello-oligosaccharide (COS), 40 mg/ml xylo-oligosaccharide (XOS) and 20 mg/ml chitin-oligosaccharide (CHOS) for 24 h. GO term enrichment is expressed as significantly different fold enrichment of mapped genes (FDR p<0.05). The dot size (and numbers beside the dots) indicates the number of significantly up-regulated genes (see Figure 2A) associated with the process and the dot color indicates the significance of the enrichment. See Supplementary Figure 1-3 for GO enrichment analysis for independent treatment of COS, XOS and CHOS.

### Assessment of Elicitor Activity of Different Lengths of Cello- and Xylo-oligosaccharide

Previous studies have shown that the recognition of oligosaccharides by plant receptors varies in its efficiency depending on their degree of polymerization (DP). In the case of chitin-oligosaccharides, the formation of the receptor complex is activated by chitin of DP8 in Arabidopsis and rice (Liu et al. 2012, Cao et al., 2014). To determine polymerization levels of COS and XOS, which are most active in inducing resistance responses in Arabidopsis, activation of immune responses was measured using Arabidopsis p*WRKY33-LUC* transformants. The different DP of oligosaccharides from the hydrolysis product of cotton linters (COS) or corn cobs (XOS) was separated by HPLC (Supplementary Figures 4 and 5) and subjected to the elicitor activity assay using Arabidopsis p*WRKY33-LUC*. Treatment with cellotriose (DP3) elicited the strongest activation, while longer DP4 and DP5 showed relatively lower elicitor activity. Cellobiose (DP2) still retained elicitor activity, but its activity was significantly lower (Figure 6A). For XOS, xylotetraose (DP4) showed the strongest elicitor activity followed by and xylotriose (DP3), while shorter DP2 and longer DP5 showed apparently lower activity (Figure 6B). These results indicated that DP3 cello-oligosaccharide and DP4 xylo-oligosaccharide are well-recognized by Arabidopsis cells for induction of disease resistance, thus it seems important to prepare the proper lengths of oligosaccharides with high activity when extracting COS and XOS from plant materials for practical use of these oligosaccharides.

**FIGURE 6.**
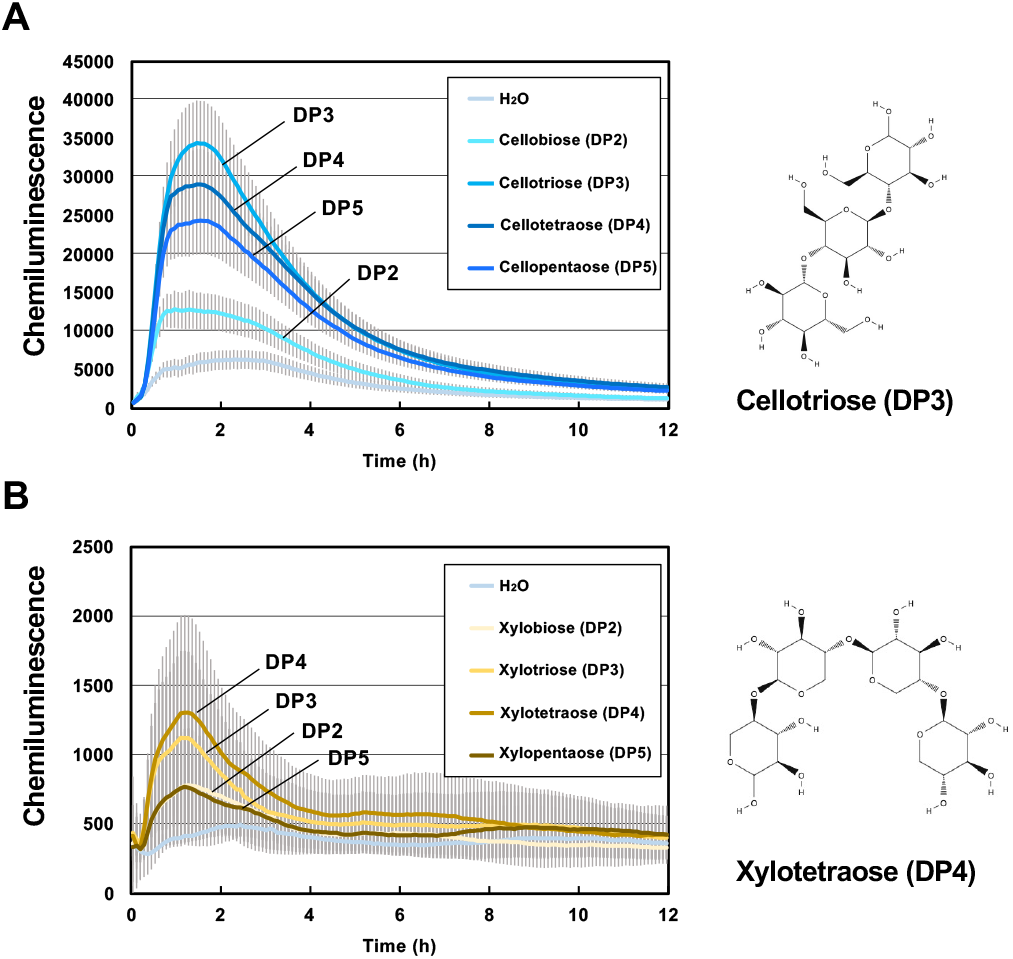
Activation of *AtWRKY33* promoter in Arabidopsis treated with different degrees of polymerization (DP) of cello-oligosaccharide and xylo-oligosaccharides. **(A)** Arabidopsis transformant pWRKY33*-*LUC was treated with 25 μg/ml cellobiose (DP2), cellotriose (DP3), cellotetraose (DP4), or cellopentaose (DP5). **(B)** Arabidopsis transformant pWRKY33*-*LUC was treated with 25 μg/ml xylobiose (DP2), xylotriose (DP3), xylotetraose (DP4), or xylopentaose (DP5). Chemiluminescence intensity was determined from an individual seedling for 12 h after treatment. Data represented as mean ± SD (n = 3) from an experiment of three repetitions.

### Pre-treatment of Oligosaccharide Mixture Induces Disease Resistance Against Powdery Mildew in Tomato

To evaluate the effect of plant resistance induced by the oligo-mix in the field, disease resistance in tomato grown in the greenhouse was evaluated. Two tomato varieties were grown in a greenhouse, sprayed with the oligo-mix once every two weeks, and naturally allowed to get infected by pathogens. Powdery mildew was observed on tomato plants after spraying for several weeks. At approx. 12 weeks (after 6 times of oligo-mix treatment), the number of colonies of powdery mildew on leaves was counted. Notably, treatment with oligo-mix significantly reduced the disease symptoms compared with the control, which was relatively susceptible to powdery mildew infection in both tested cultivars (Figure 7).

**FIGURE 7.**
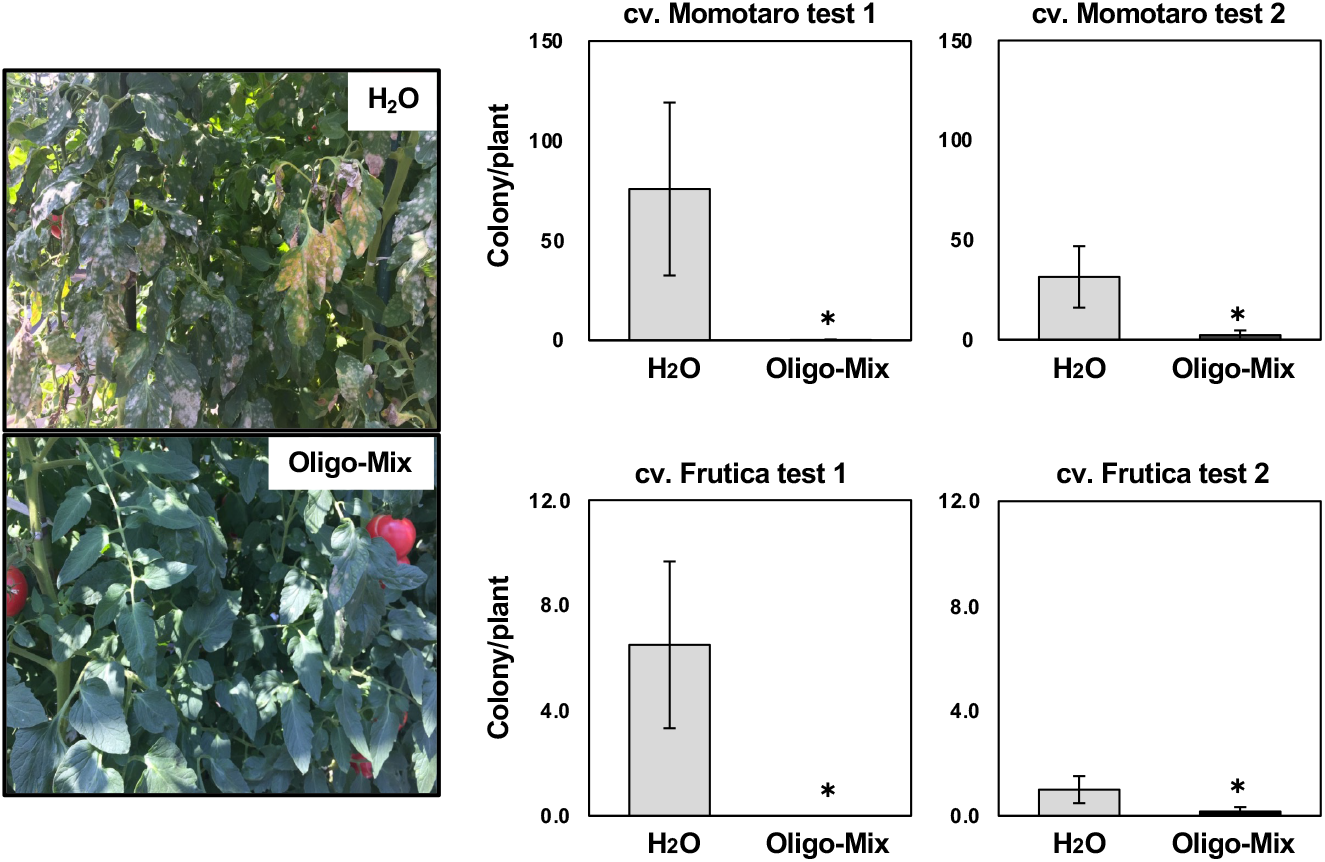
Treatment with oligo-mix significantly reduces disease symptoms of powdery mildew in tomato. Tomato plants were grown in a greenhouse and sprayed with Oligo-mix (20 mg/ml COS, 40 mg/ml XOS and 20 mg/ml CHOS) once every two weeks after 5 days of germination (6 times). Number of colonies per plant on tomato cultivars Momotaro or Frutica was counted 12 weeks after the initial treatment. Data are shown as mean ± SD of colony number per plant (n = 6). Asterisks indicate a significant difference from water-treated control as assessed by two-tailed Student’s *t*-test. *p< 0.05.

### RNAseq Analysis of Tomato Treated with Oligo-Mix

To further elucidate the effect of oligo-mix treatment on tomato, tomato plants sprayed with oligo-mix (once every week after germination, 4 times overall) were subjected to the RNAseq analysis (Supplementary Table 1). Interestingly, treatment of oligo-mix showed a tendency to increase the growth of both shoot and root (Supplementary Figure 6). Upregulated genes in tomato treated with oligo-mix, includes genes potentially involved in the plant defense against pathogens and insects, including plant basic secretory protein NtPRp27-like (Solyc01g080790, Xie and Goodwin, 2009), sugar transporter SWEET (Solyc03g097570, Chen et al., 2015), calcineurin subunit B-like (Solyc07g005900, Kurusu et al., 2010), protease inhibitor I (Solyc09g084460, Zhu-Salzman and Zeng, 2015) and some pathogenesis-related proteins (Solyc11g072830, Solyc04g076260, Solyc02g065470, van Loon et al., 2006). Genes related to the activation of ethylene production, ethylene-responsive transcription factor (Solyc01g090340) and ACC oxidase (Solyc09g089750), can be considered as defense-related since ethylene is involved in the induction of phytoalexin in Solanaceae plants (Shibata et al., 2010; Takemoto et al., 2018). Genes inducible by oligo-mix also include a group of genes presumed to be involved in the construction of the cell wall including UDP-glycosyltransferase 76E1 (Solyc10g085280, Piršelová and Matušíková, 2013), xyloglucan endotransglucosylase (Solyc11g066270, Ishida and Yokoyama, 2022) and caffeoyl-CoA O-methyltransferase (Solyc09g082660, Zhong et al., 1998). Notably, oligo-mix-induced genes includes many genes related to chloroplasts, such as chloroplastic ribosomal proteins (e.g. Solyc00g500144), TIC 214 (e.g. Solyc00g500279), photosystem I assembly protein Ycf3 (e.g. Solyc00g500058), ATP synthase subunit α (Solyc00g500358) and Photosystem I P700 (Solyc00g500199) (Supplementary Table 1). These results indicate that oligo-mix have the potential to activate plant disease resistance and growth in tomato.

### Concluding Remarks

The plant cell wall is the first line of defense between plant and microbial pathogens. It is composed of complex structures of carbohydrate polysaccharides as well as proteins that protect plants from infection (Somerville et al., 2004; Cosgrove, 2022). Pathogens have arsenals of cell wall degradation enzymes to digest cell walls such as pectate lyase, cellulases, xylanases, and proteases, resulting in breakdown molecules released upon plant-microbe interactions. At the site of infection, plants infected by pathogens simultaneously receive multiple PAMPs and DAMPs and use the combined signals to activate an appropriate resistance response and recovery from the damage.

Generally, enhancement of the resistance response is a trade-off for growth, for example, excessive use of resistance inducers such as SA analogs is known to cause negative effect on growth (Tripathi et al., 2019). In the short term, oligosaccharide elicitors are resistance inducers, as shown by the RNAseq analysis of oligosaccharide elicitor-treated Arabidopsis, where genes induced at 24 h after treatment were mainly related to resistance responses (Figures 3-5, Supplementary Figures 1-3). Meanwhile, when tomatoes were treated with oligosaccharide elicitors on a regular basis over a longer period, the induction of genes associated with resistance was less clear than in Arabidopsis, while the expression of a group of genes associated with chloroplasts was induced (Supplementary Table 1). In fact, oligosaccharide treatment did not inhibit tomato growth, but tended to promote stem and root growth (Supplementary Figure 6). The growth-promoting effect of oligosaccharide elicitors in tomato may have resulted from the long-term effect of the reported growth-promoting effect of XOS (Cutillas-Iturralde and Lorences, 1997; Kaida et al., 2010; Claverie et al., 2018), which could have contributed to the favorable basic conditions of the plants, such as enhanced root growth. In order to create an effective biostimulant material that shows beneficial effects on plants for both defense against pathogen and growth, mixing a greater variety of PAMPs and DAMPs may be effective. Appropriate blending of substances other than the 3 elicitors used in this study, such as oligogalacturonides and arabinoxylan-oligosaccharides (Voxeur et al., 2019; Mélida et al., 2020), could lead to the development of more effective biostimulants.

## Author contributions

MS and DT designed the research. PS, H Kato, SI, H Kimoto, PC, AS, H Kobayashi, TS and DT conducted the experiments. PS, H Kato, SI, MC, AT, H Kimoto and DT analyzed data, SI, MC, AT, AF, RT, IS, SC, KK and DT supervised the experiments. PS, H Kato, MC, H Kimoto, MS and DT contributed to the discussion and interpretation of the results. PS and DT wrote the manuscript. MC and DT edited the manuscript.

## Funding

This work was supported partially by a Grant-in-Aid for Scientific Research (B) (20H02985 and 23H02212) to DT from the Japan Society for the Promotion of Science, and donation from Resonac Corporation (former Showa Denko K.K., Tokyo, Japan).

## Conflict of interest

This study was partially funded by Resonac corporation (Japan). MS is an employee of Resonac corporation.

## Acknowledgments

We thank Prof. Kazuhito Kawakita (Nagoya University, Japan) for valuable suggestions. We are also grateful to Mr. Masahiro Maesaka and Ms. Yuki Iwamoto (Nagoya University), and Ms. Ayami Furuta (Chubu University, Japan) for their technical support. We would like to acknowledge the Japanese Ministry of Education, Culture, Sports and Technology (MEXT) for supporting Pring Sreynich to pursue study in Japan on scholarship.

## SUPPLEMENTAL DATA

**SUPPLEMENTARY FIGURE 1** | Gene ontology (GO) enrichment analysis of up-regulated genes in Arabidopsis treated with 20 mg/ml cello-oligosaccharide (COS) for 24 h. GO term enrichment is expressed as significantly different fold enrichment of mapped genes (FDR p<0.05). The dot size (and numbers beside the dots) indicates the number of significantly up-regulated genes (see Figure 2A) associated with the process and the dot color indicates the significance of the enrichment.

**SUPPLEMENTARY FIGURE 2** | Gene ontology (GO) enrichment analysis of up-regulated in Arabidopsis treated with 40 mg/ml xylo-oligosaccharide (XOS) for 24 h. GO term enrigeneschment is expressed as significantly different fold enrichment of mapped genes (FDR p<0.05). The dot size (and numbers beside the dots) indicates the number of significantly up-regulated genes (see Figure 2A) associated with the process and the dot color indicates the significance of the enrichment.

**SUPPLEMENTARY FIGURE 3** | Gene ontology (GO) enrichment analysis of up-regulated in Arabidopsis treated with 20 mg/ml chitin-oligosaccharide (CHOS) for 24 h. GO term enrigeneschment is expressed as significantly different fold enrichment of mapped genes (FDR p<0.05). The dot size (and numbers beside the dots) indicates the number of significantly up-regulated genes (see Figure 2A) associated with the process and the dot color indicates the significance of the enrichment.

**SUPPLEMENTARY FIGURE 4** | Different lengths of cello-oligosaccharides were separated and collected by HPLC equipped with a refractive index detector (Shimadzu RID 10-ATVP) and a fraction collector (Shimadzu FRC 10A). See materials and method for detail.

**SUPPLEMENTARY FIGURE 5** | Different lengths of xylo-oligosaccharides were separated and collected by HPLC equipped with a refractive index detector (Shimadzu RID 10-ATVP) and a fraction collector (Shimadzu FRC 10A). See materials and method for detail.

**SUPPLEMENTARY FIGURE 6** | Oligo-mix could promote growth of tomato plant.

Effect of oligo-mix on the growth of tomato plants. Tomato (cv. Renaissance) was treated with water or oligo-mix (20 mg/ml COS, 40 mg/ml XOS and 20 mg/ml CHOS) from 3 days after the seedling germination, once every week (total 4 times). One week after the last treatment, growth (fresh and dry weight) of tomato was measured, and the leaf samples were used for the extraction of total RNA for RNAseq analysis (Supplementary Table 1). Data are means ± standard error (n = 4).

**SUPPLEMENTARY TABLE 1** | Tomato genes upregulated by oligo-mix treatment.

