## Supplemental Data for "Induction of plant disease resistance by mixed oligosaccharide elicitors prepared from plant cell wall and crustacean shells"

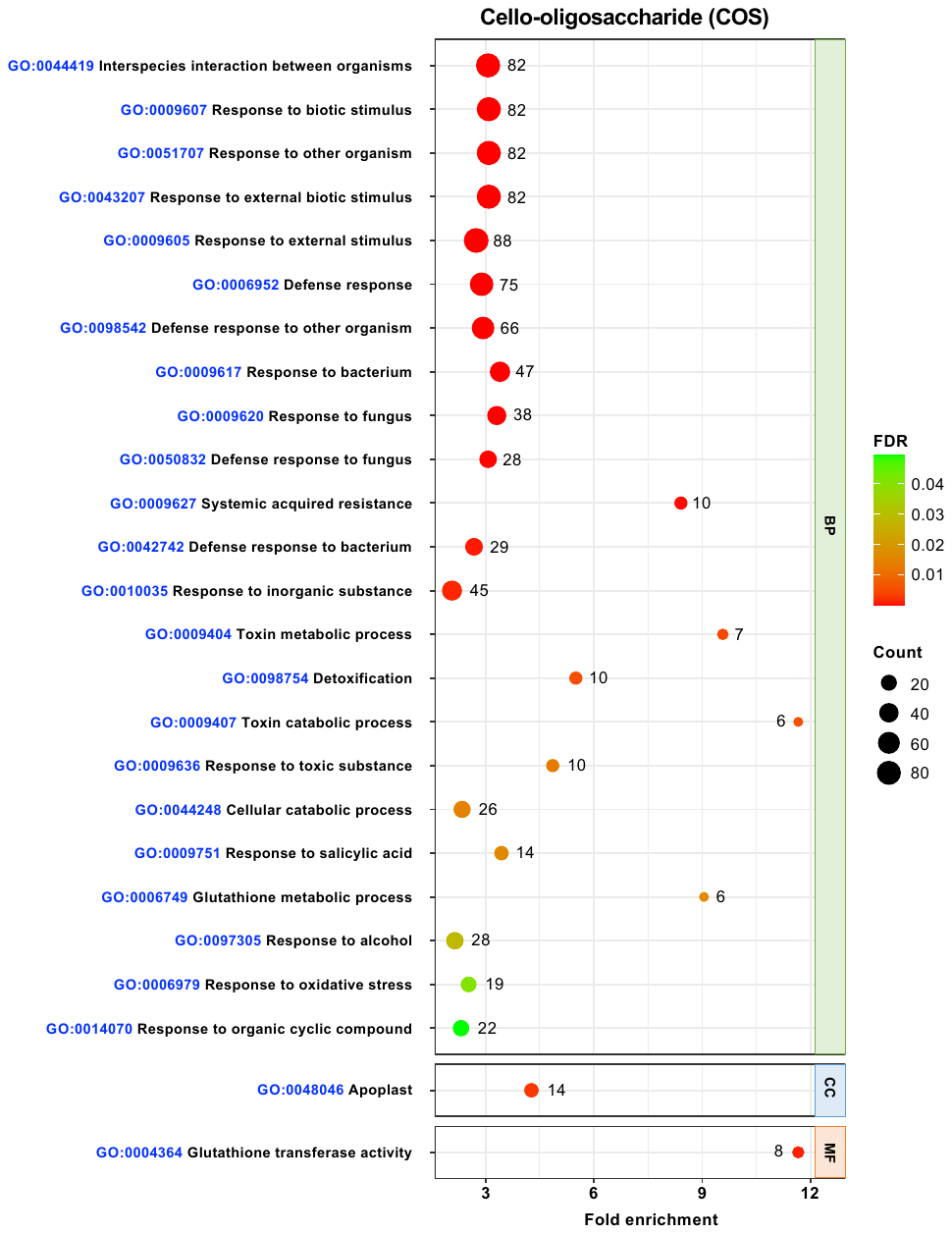

**SUPPLEMENTARY FIGURE 1 |** Gene ontology (GO) enrichment analysis of up-regulated genes in Arabidopsis treated with 20 mg/ml cello-oligosaccharide (COS) for 24 h. GO term enrichment is expressed as significantly different fold enrichment of mapped genes (FDR p<0.05). The dot size (and numbers beside the dots) indicates the number of significantly up-regulated genes (see Figure 2A) associated with the process and the dot color indicates the significance of the enrichment.

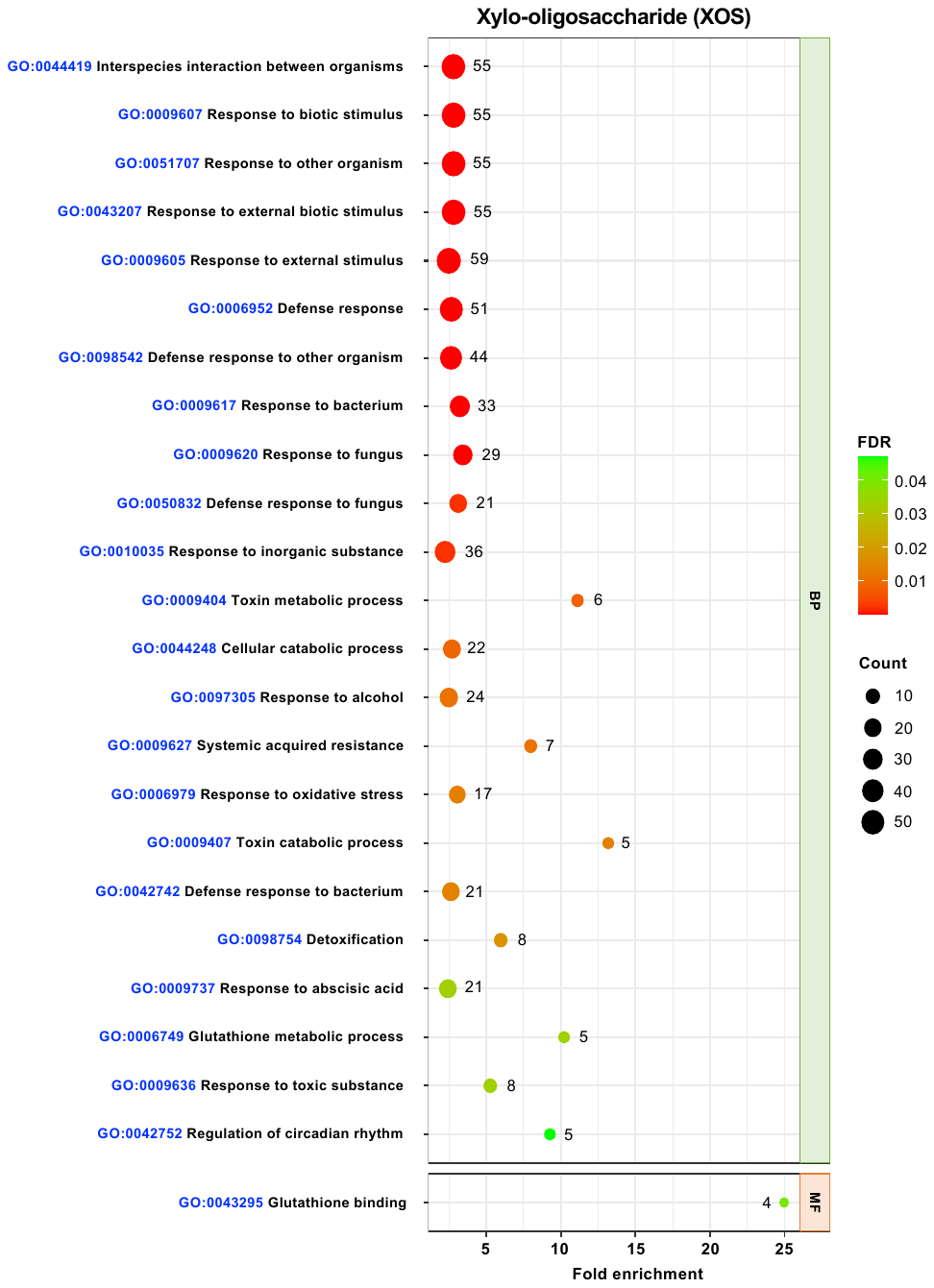

**SUPPLEMENTARY FIGURE 2** | Gene ontology (GO) enrichment analysis of up-regulated in Arabidopsis treated with 40 mg/ml xylo-oligosaccharide (XOS) for 24 h. GO term enrigeneschment is expressed as significantly different fold enrichment of mapped genes (FDR p<0.05). The dot size (and numbers beside the dots) indicates the number of significantly up-regulated genes (see Figure 2A) associated with the process and the dot color indicates the significance of the enrichment.

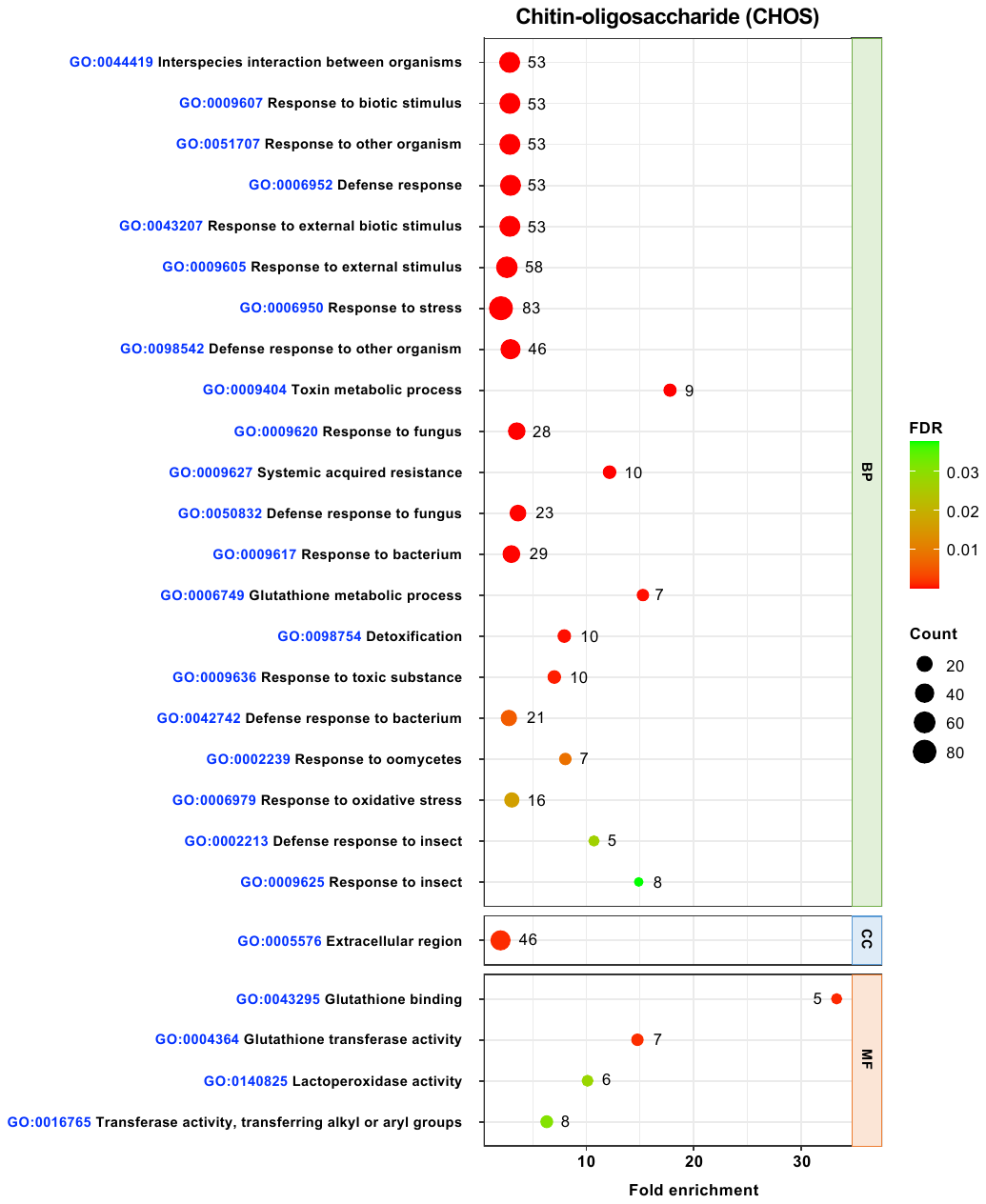

**SUPPLEMENTARY FIGURE 3 |** Gene ontology (GO) enrichment analysis of up-regulated in Arabidopsis treated with 20 mg/ml chitin-oligosaccharide (CHOS) for 24 h. GO term enrigeneschment is expressed as significantly different fold enrichment of mapped genes (FDR p<0.05). The dot size (and numbers beside the dots) indicates the number of significantly up-regulated genes (see Figure 2A) associated with the process and the dot color indicates the significance of the enrichment.

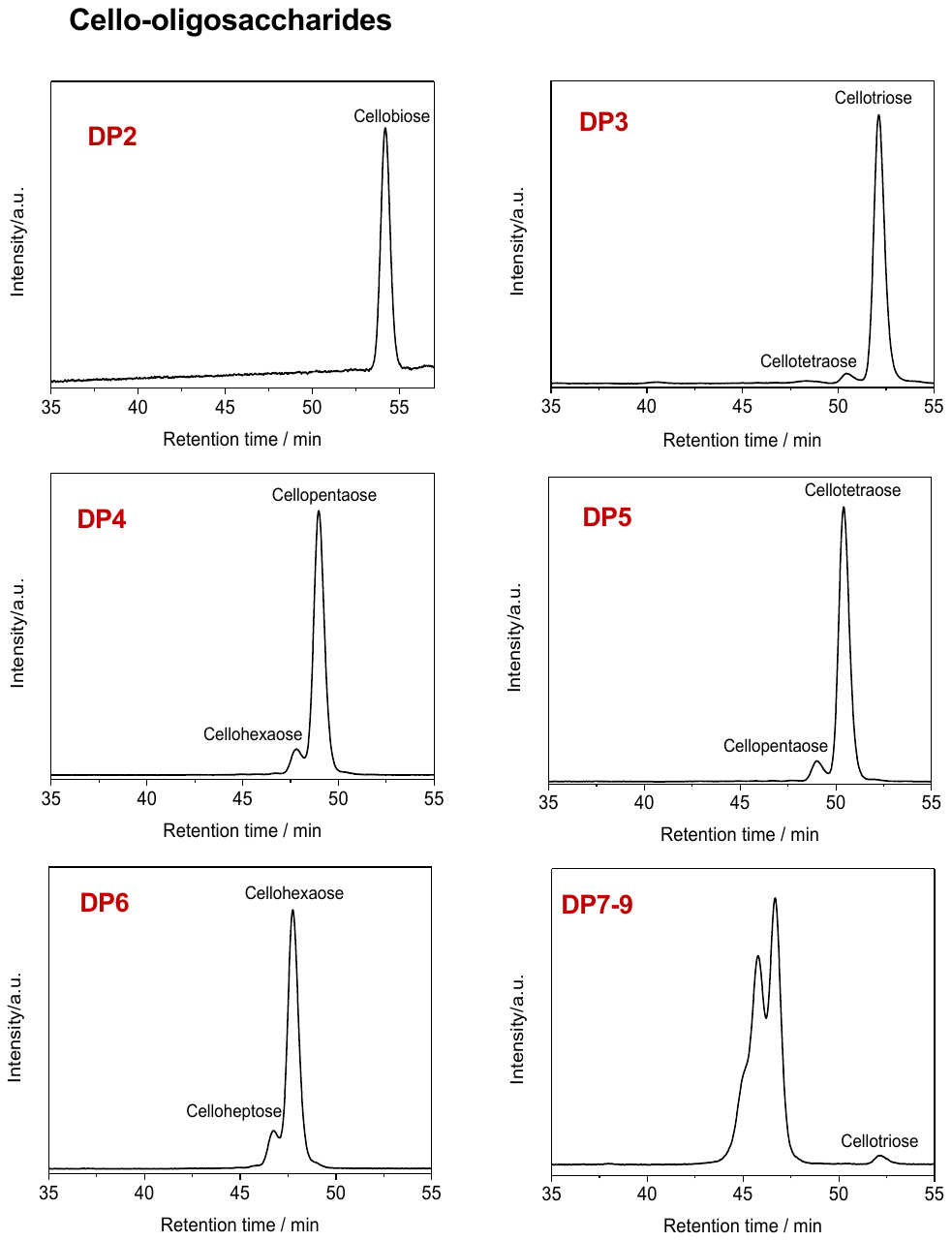

**SUPPLEMENTARY FIGURE 4 |** Different lengths of cello-oligosaccharides were separated and collected by HPLC equipped with a refractive index detector (Shimadzu RID 10-ATVP) and a fraction collector (Shimadzu FRC 10A). See materials and method for detail.

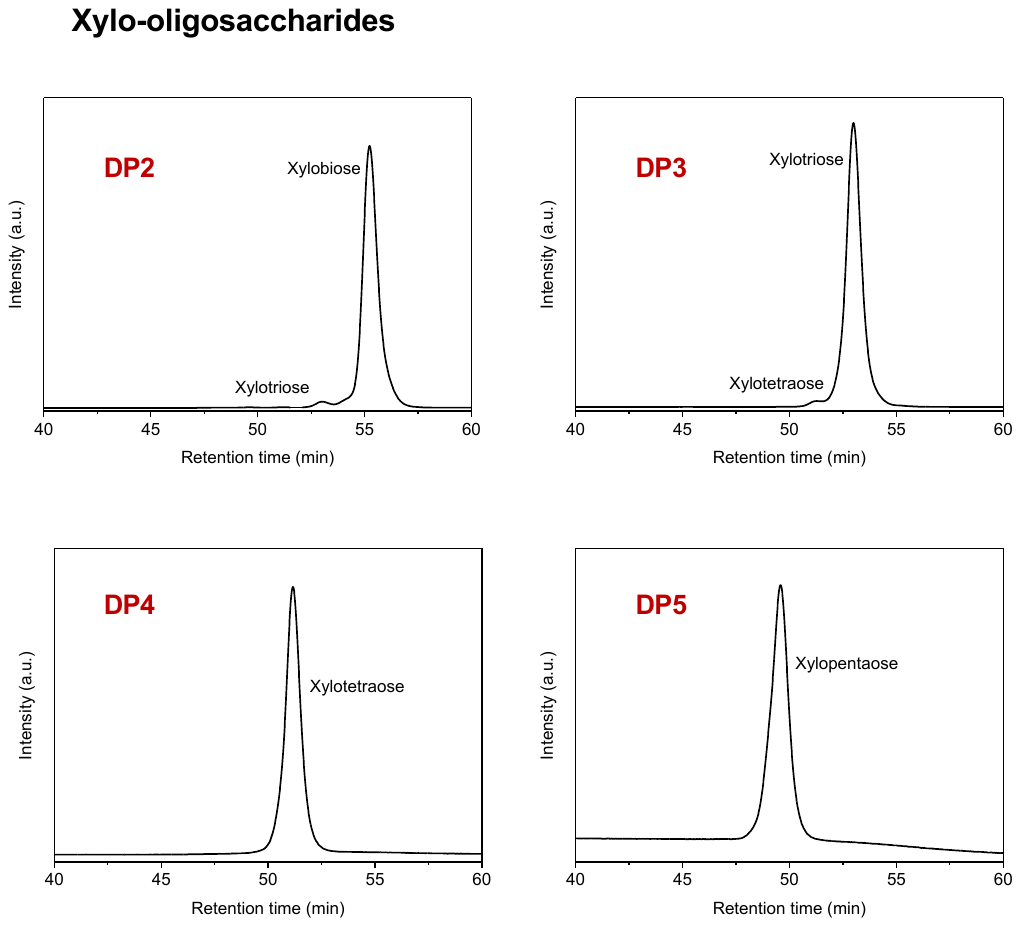

**SUPPLEMENTARY FIGURE 5 |** Different lengths of xylo- oligosaccharides were separated and collected by HPLC equipped with a refractive index detector (Shimadzu RID 10-ATVP) and a fraction collector (Shimadzu FRC 10A). See materials and method for detail.

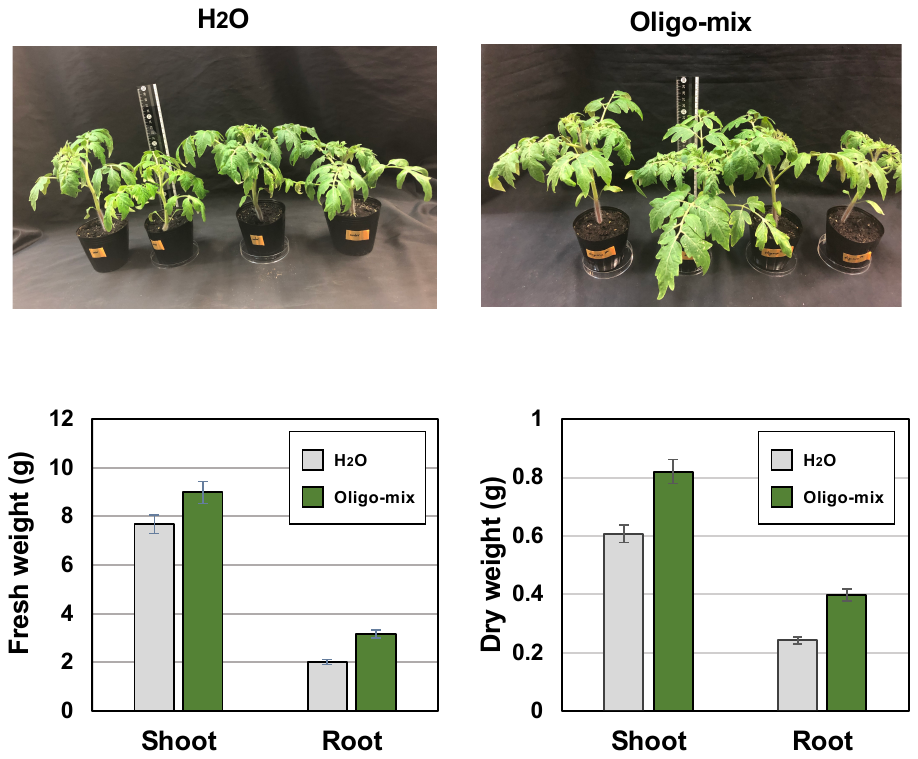

**SUPPLEMENTARY FIGURE 6 |** Oligo-mix could promote growth of tomato plant.

Effect of oligo-mix on the growth of tomato plants. Tomato (cv. Renaissance) was treated with water or oligo-mix (20 mg/ml COS, 40 mg/ml XOS and 20 mg/ml CHOS) from 3 days after the seedling germination, once every week (total 4 times). One week after the last treatment, growth (fresh and dry weight) of tomato was measured, and the leaf samples were used for the extraction of total RNA for RNAseq analysis (Supplementary Table 1). Data are means ± standard error (n = 4).

**]**

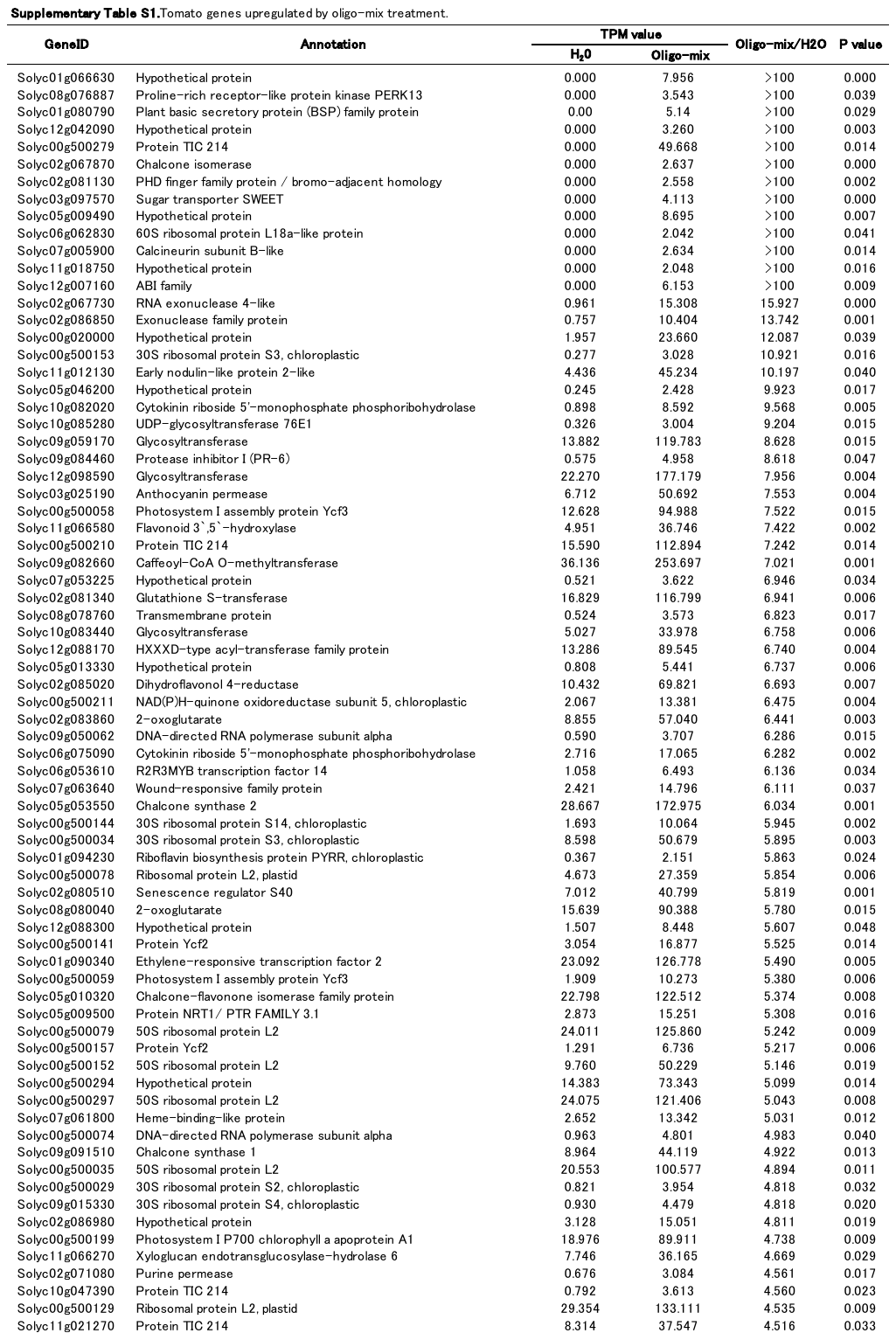

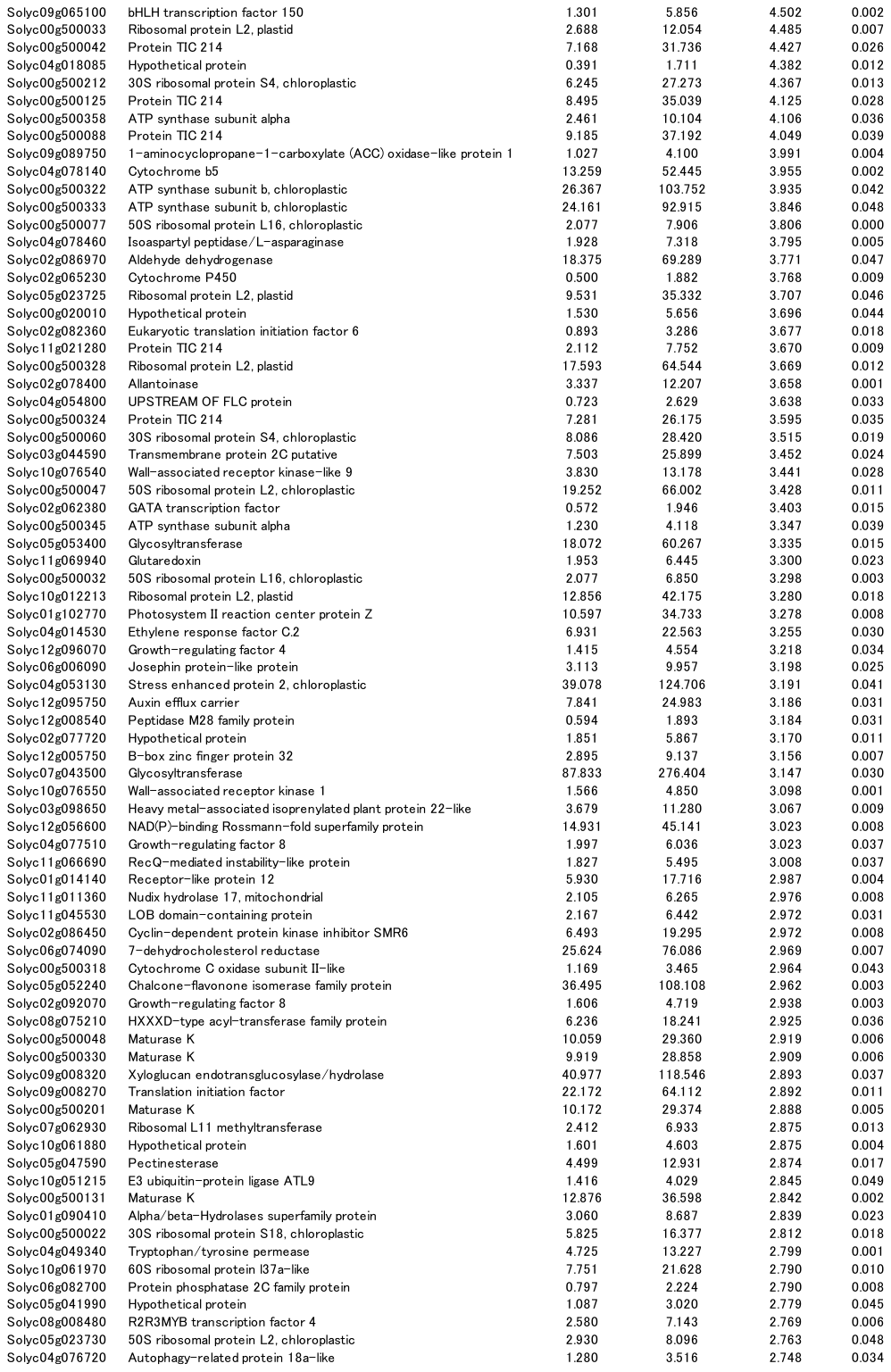

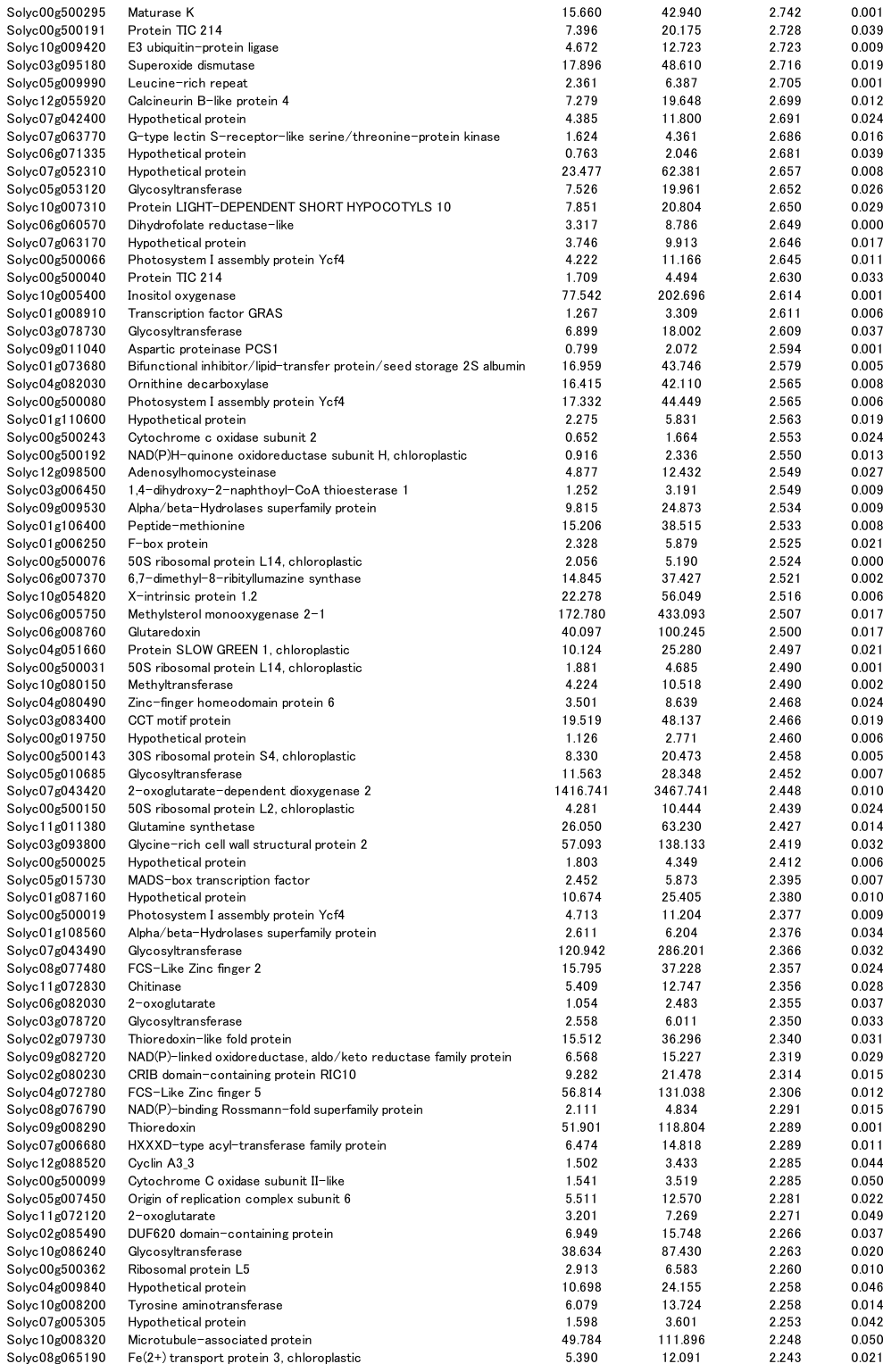

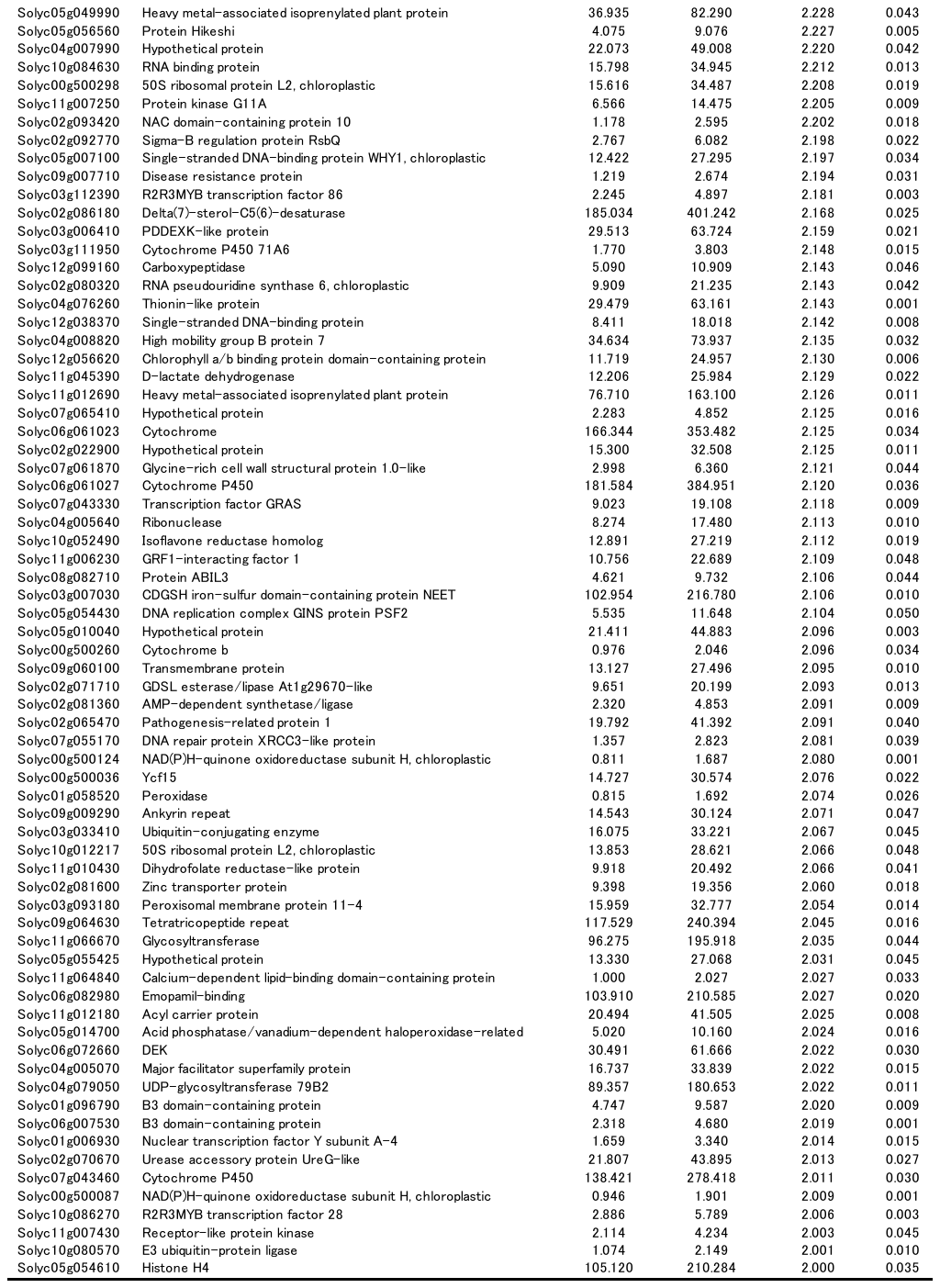
